# Cut-and-Paste DNA Insertion with Engineered Type V-K CRISPR-associated Transposases

**DOI:** 10.1101/2022.01.07.475005

**Authors:** Connor J. Tou, Benno Orr, Benjamin P. Kleinstiver

## Abstract

CRISPR-associated transposases (CASTs) enable recombination-independent, multi-kilobase DNA insertions at RNA-programmed genomic locations. Type V-K CASTs offer distinct technological advantages over type I CASTs given their smaller coding size, fewer components, and unidirectional insertions. However, the utility of type V-K CASTs is hindered by a replicative transposition mechanism that results in a mixture of desired simple cargo insertions and undesired plasmid co-integrate products. Here, we overcome this limitation by engineering new CASTs with dramatically improved product purity. To do so, we compensate for the absence of the TnsA subunit in multiple type V-K CASTs by engineering a Homing Endonuclease-assisted Large-sequence Integrating CAST compleX, or HELIX system. HELIX utilizes a nicking homing endonuclease (nHE) fused to TnsB to restore the 5 “ nicking capability needed for dual-nicking of the DNA donor. By leveraging distinct features of both type V-K and type I systems, HELIX enables cut-and-paste DNA insertion with up to 99.3% simple insertion product purity, while retaining robust integration efficiencies on genomic targets. Furthermore, we demonstrate the versatility of this approach by generating HELIX systems for other CAST orthologs. We also establish the feasibility of creating a minimal, 3-component HELIX, simplifying the number of proteins that must be expressed. Together, HELIX streamlines and improves the application of CRISPR-based transposition technologies, eliminating barriers for efficient and specific RNA-guided DNA insertions.

## Main

Programmable insertion of multi-kilobase DNA sequences into genomes without reliance on homologous recombination and double-stranded breaks (DSBs) would offer new capabilities for precision genome editing. Methods for genomic integration typically rely on viral vectors^1,2^ or transposons^3-7^, both of which lack programmability and thus insert stochastically throughout the genome, or nucleases coupled with DNA donors^8-10^ that rely on cytotoxic DSBs and host homologous recombination machinery. Additionally, recombineering systems in bacteria are low efficiency^11^ without cointegration of a selectable marker^12^ or CRISPR-Cas counterselection^13^. CRISPR-associated transposases (CASTs) are a promising new approach for programmable, recombination-independent DNA insertions through an interplay between transposase machinery and CRISPR effector(s) to direct RNA-guided transposition^14-16^.

The two main classes of CASTs, types I and V-K, have distinct and complementary properties. While characterized type I CASTs exhibit high on-target specificity and generally only result in the intended simple insertion gene products^17^ (though with exceptions^18^), the larger number of Cas genes, stoichiometric complexity, and coding size may limit downstream tool development in other organisms such as eukaryotic cells. Additionally, the tendency of some type I systems to result in bidirectional insertions leads to undesirable edit impurity^15^ (**Fig. 1a**). In comparison, type V-K CASTs are more compact in terms of coding size, contain only four core components, and deliver complete or near-complete unidirectional insertions. However, type V-K CASTs lead to a problematic mixture of simple insertion and co-integrate gene products, the latter of which consists of cargo duplication and full plasmid backbone insertion^4,6^ (**Fig. 1b**), and are generally lower specificity^14,16,17^. For genome editing applications, an ideal DNA insertion technology would enable programmable, unidirectional, recombination-independent, and pure simple insertion gene products with minimal components and minimally sized machinery. Therefore, we sought to develop an engineered CAST technology combining the simplicity and orientation predictability of type V-K systems with the product purity of type I systems.

**Figure 1.**
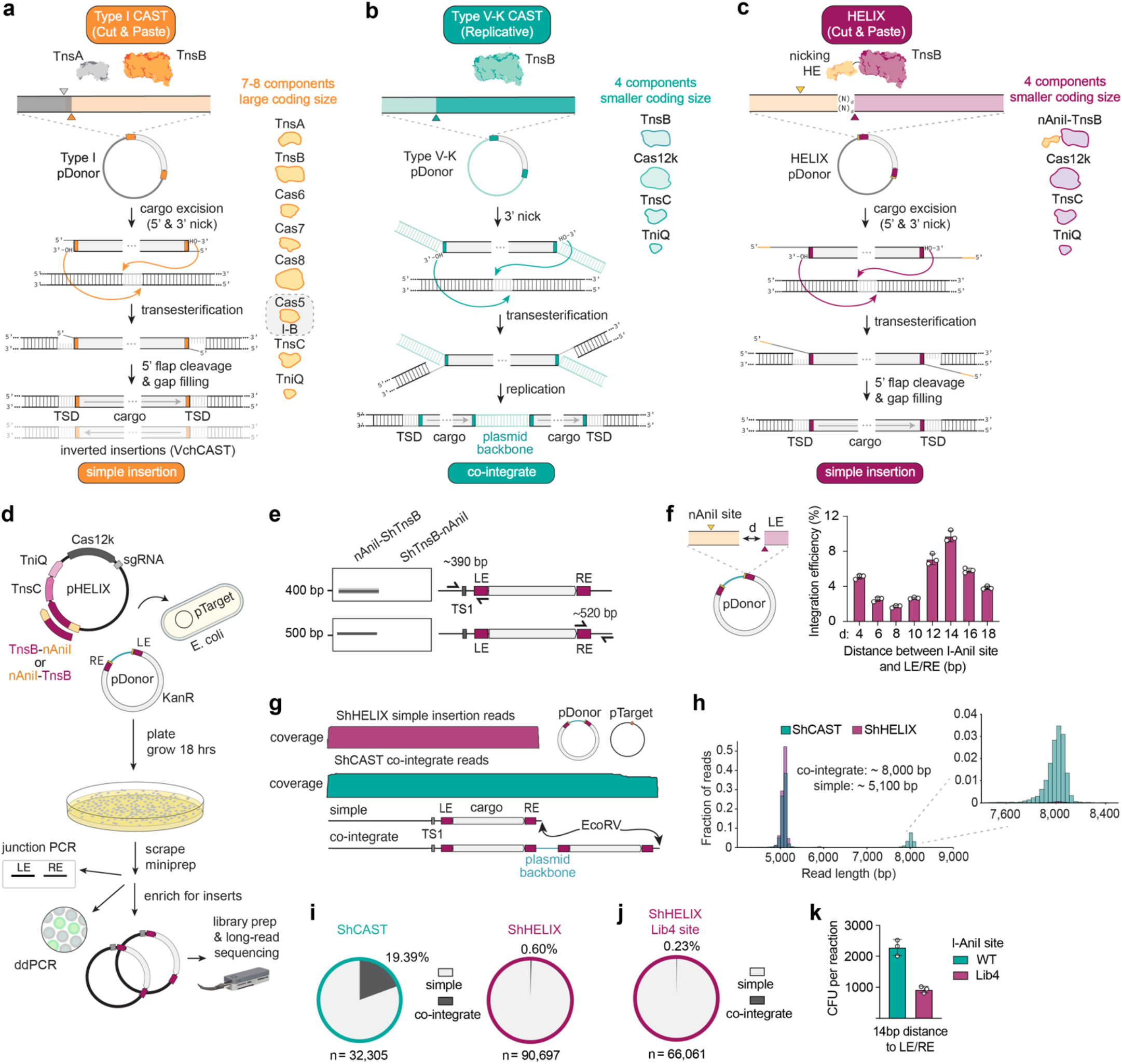
Development and characterization of HELIX. **a**-**c**, Schematics of type I and type V-K CASTs and HELIX (**panels a**-**c**, respectively) and their transposition mechanisms that result in simple insertion or co-integrate gene products. **d**, Workflow for transposition experiments targeting plasmid substrates. **e**, Transposition assessed via junction PCRs across the LE/RE at TS1 in pTarget. Experiments were performed with nAniI fused to the N- or C-terminus of TnsB when using pDonor without I-AniI sites. **f**, Quantification of DNA integration efficiency on plasmids when using ShHELIX and a donor plasmid with a range of distances (d) between the I-AniI site and LE/RE, assessed via ddPCR using miniprepped DNA. **g**, Coverage of expected insertion products into pTarget from long-read sequencing using a subset of exemplary simple insertion reads for ShHELIX and co-integrate reads for ShCAST (coverage from ShHELIX co-integrate reads and ShCAST simple insertion reads omitted for simplicity). **h**, Read length distribution when using ShCAST and ShHELIX with a sgRNA targeting TS1 on pTarget from long-read sequencing data. The top right panel is a zoomed-in image of the ∼8,000 bp read-length peak. **i**, Comparison of simple insertion and co-integrate product proportions of transposed products forShCAST and ShHELIX constructs when using a pDonor with I-AniI sites 14 bp from the LE/RE and oriented to confer a 5 “ nick, assessed via long-read sequencing. **j**,**k**, Transposition product purity (**panel j**) and CFUs (**panel k**) when using a Lib4 I-AniI site on pDonor (with a distance of 14 bp between the Lib4 sites and the LE/RE), which was previously shown to increase affinity of wild type I-AniI by 5-fold. For **panels f** and **k**, mean, SD, and individual data points shown for n = 3. TSD, target-site duplication; LE and RE, left and right transposon ends, respectively; sgRNA, single guide RNA; ddPCR, droplet digital PCR.

Type I and type V-K CASTs differ by the presence or absence of the TnsA subunit, respectively, one major distinction defining their simple insertion to co-integrate product purity. In both Tn7 transposons and type I CASTs, TnsA and TnsB carry out the 5 “ and 3 “ donor nicking reactions, respectively, resulting in simple insertions via cut-and-paste transposition (**Fig. 1a**). In Tn5053 transposons and type V-K CASTs, which lack TnsA, and also in Tn7 transposons and modified type I systems with catalytically dead TnsA^17,20^, only 3 “ donor nicking occurs via TnsB. Singly-nicked donors result in a substantial fraction of co-integrate insertions through replicative, instead of cut-and-paste, transposition^21^ (**Fig. 1b**). Therefore, to create a co-integrateless type V-K CAST, we hypothesized that restoring the absent function of TnsA by fusing an orthogonal DNA nickase to TnsB would enable cut-and-paste transposition via 5 “ and 3 “ donor nicking (**Fig. 1c**). An ideal nickase would be small (to add minimal coding size to the system), have a predictable nicking site and strand preference, have high specificity to prevent unintended on- or off-target nicking, and would function in various organisms for downstream tool development and application. Potential options include nicking restriction endonucleases^22^, nicking Cas variants^9,23,24^, the catalytic portion or full TnsA enzyme from type I CASTs or Tn7 transposons^25^, or phage HNH endonucleases^26^. However, each comes with drawbacks regarding size, sequence and strand specificity, and/or required protein-protein interactions.

One family of enzymes that fits these criteria are homing endonucleases (HEs), which are small nucleases that can generate DSBs with relatively high specificity. LAGLIDADG HEs (LHE) have been harnessed for genome editing in bacterial and human cells because of their specificity and their moderate reprogrammability via protein engineering or chimeric assembly^27^. The LHE from *Aspergillus nidulans* (I- AniI) has a small coding sequence (254 amino acids), cleaves a 19-bp asymmetric DNA target sequence, and has been previously engineered to be a sequence-specific nickase through a single K227M mutation^28^ (nAniI). Furthermore, a hyperactive variant of I-AniI, termed Y2 I-AniI, has been shown to have a 9-fold higher affinity for its cognate target site^29^. Therefore, we hypothesized that either nAniI or Y2 nAniI could be fused to TnsB to enable dual nicking on the donor plasmid required for cut-and-paste DNA insertion with type V-K CASTs (**Fig. 1c**). Together, the nHE fusion to TnsB along with the remaining CAST components form an HE-assisted Large-sequence Integrating CAST compleX, or HELIX.

## Results

### Development and optimization of HELIX

We first determined whether nAniI could adequately substitute for the lack of TnsA in the canonical type V-K transposon from *Scytonema hofmannii* (ShCAST)^14^. To do so, we constructed a series of ShHELIX expression plasmids that each contained: (1) a single guide RNA (sgRNA) targeting target site 1 (TS1) on a separate target plasmid (pTarget), (2) Cas12k, (3) TniQ, (4) TnsC, and (5) nAniI fused to the N- or C-terminus of TnsB (**Fig. 1d**). ShCAST or ShHELIX expression plasmids were co-transformed with a donor plasmid (pDonor, containing a 2.1kb cargo) with left and right transposon ends (LE and RE, respectively) into an *E. coli* strain harboring pTarget (**Fig. 1d**). To determine whether transposition occurred when using the nAniI fusions to TnsB, we performed junction PCR across both the LE and RE within pTarget on miniprepped DNA from scraped colonies. Interestingly, fusion of nAniI to the N-terminus of TnsB supported RNA-guided DNA insertion (**Fig. 1e**). C-terminal fusions of nAniI to TnsB did not result in transposition, suggesting that the C-terminal TnsC interacting domain of TnsB is less accommodating to fusion proteins^30^.

Next, we assembled eight additional donor plasmids harboring variable distances between the LE/RE and the I-AniI target site (oriented to confer a 5 “ nick on pDonor) (**Fig. 1f**). When co-transforming ShCAST or ShHELIX (with N-terminal nAniI-TnsB fusion) expression plasmids along with pDonor into our pTarget strain, we observed similar numbers of transformant colonies when using ShCAST and ShHELIX constructs, indicating no cell-viability defect due to HELIX (**Sup. Fig. 1a**). We assessed integration efficiency via droplet digital PCR (ddPCR), and with ShHELIX we observed a range of integration efficiencies across different spacings on pDonor, with 14 bp spacing yielding the highest integration (**Fig. 1f**). Surprisingly, ShCAST also exhibited variable integration efficiency depending on the spacing between the I-AniI site and LE/RE (where, unlike with ShHELIX, the I-AniI site has no role in transposition). Spacings of 4-12 bp on pDonor resulted in substantially higher insertion efficiencies than a pDonor without I-AniI sites (**Sup. Fig. 1b**). Altering the position of the I-AniI site modifies the sequence directly adjacent to the LE/RE on pDonor, suggesting that the composition of the flanking sequence, particularly the first 12 bp, may be an important determinant of integration efficiency (**Sup. Fig. 1b**). Separately, we also performed integration experiments using Y2 nAniI fused to TnsB (Y2 ShHELIX) and observed substantially fewer colonies, with peak numbers using 14 bp spacing (**Sup. Fig. 3a** and **Sup Note. 1**). For subsequent experiments, HELIX constructs with nAniI-TnsB fusions, and pDonors with 14 bp between the I-AniI sites and LE/RE, were used.

Next, we utilized long-read sequencing to assess whether restoration of the 5 “ nick on pDonor with ShHELIX could improve product purity compared to canonical ShCAST. We enriched for transposed products from our miniprepped plasmid pool (**Sup. Fig. 2**), linearized enriched plasmid DNA, and performed long-read sequencing to determine the proportion of simple insertions to co-integrates (**Figs. 1g -1i**). With ShCAST, we observed 19.39% co-integrates, consistent with previous results^6^ (**Fig. 1i**). Strikingly, ShHELIX nearly eliminated co-integrates, resulting in a reduction to only 0.6% of all products (a 32-fold decrease when compared to ShCAST; **Figs. 1h** and **1i**). Additionally, we did not observe I-AniI sites in insertion product reads, suggesting that the 5 “ flap harboring these sequences is removed during resolution of HELIX-mediated transposition. We also performed long-read sequencing with Y2 ShHELIX and similarly observed a stark increase in simple insertion product purity (**Sup. Figs. 3b-d**).

We also performed a series of control experiments to further characterize ShHELIX (**Sup. Note 2**). First, a catalytically attenuated variant of I-AniI (K227M, Q171K) decreased co-integrates 1.8-fold compared to ShCAST (presumably due to incomplete inactivation of I-AniI nicking) (**Sup. Fig. 4a**). Secondly, a pDonor lacking an I-AniI target site resulted in a 1.7-fold reduction in co-integrates compared to ShCAST (**Sup. Fig. 4a**). Next, experiments using a pDonor with a “flipped” I-AniI site that places the nick on the same strand as the TnsB nick resulted in a 10-fold decrease in co-integrates (**Sup. Fig. 4b**). Hairpin intermediate formation or lesion-induced Shapiro intermediate cleavage (in addition to the possibility of low-level DSB-mediated cargo excision), may be occurring to result in simple insertion products (**Sup. Fig. 4c**). Finally, use of a “Lib4” I-AniI site on pDonor-14, found previously to increase the affinity of wild type I-AniI by 5-fold^31^, further decreased co-integrates relative to ShHELIX to 0.23% of all transposition products (for an 84-fold decrease in co-integrates compared to ShCAST) (**Fig. 1j**). However, this product purity improvement was also accompanied by a reduction in CFUs (**Sup. Note 1** and **Fig. 1k**). Altogether, ShHELIX coupled with an I-AniI site oriented on pDonor to confer a 5 “ nick demonstrates the most prominent increase in simple insertion to co-integrate percentage, leading to near-perfect product purity on a plasmid target.

### Characterization of HELIX on genomic targets

Encouraged by our transposition results on plasmid targets, we then explored the efficacy of ShHELIX- mediated DNA integration at genomic sites. We performed transformations using similar constructs to the plasmid targeting experiments but instead with sgRNAs targeting the genome and without pTarget (**Fig. 2a**). First, we tested the effect of two different lengths of amino acid linkers between nAniI and TnsB on genomic integration efficiency across our set of eight donor plasmids containing varying distances between the I-AniI sites and the LE/RE. Experiments were performed with a previously characterized sgRNA^14^ against a genomic target site (TS2). For both amino acid linkers, we observed the highest integration efficiency with a 14 bp spacing between the I-AniI site and LE/RE (**Fig. 2b**), which aligned with our plasmid targeting results. All detectable insertions were in the T-LR orientation (**Fig. 2c**).

**Figure 2.**
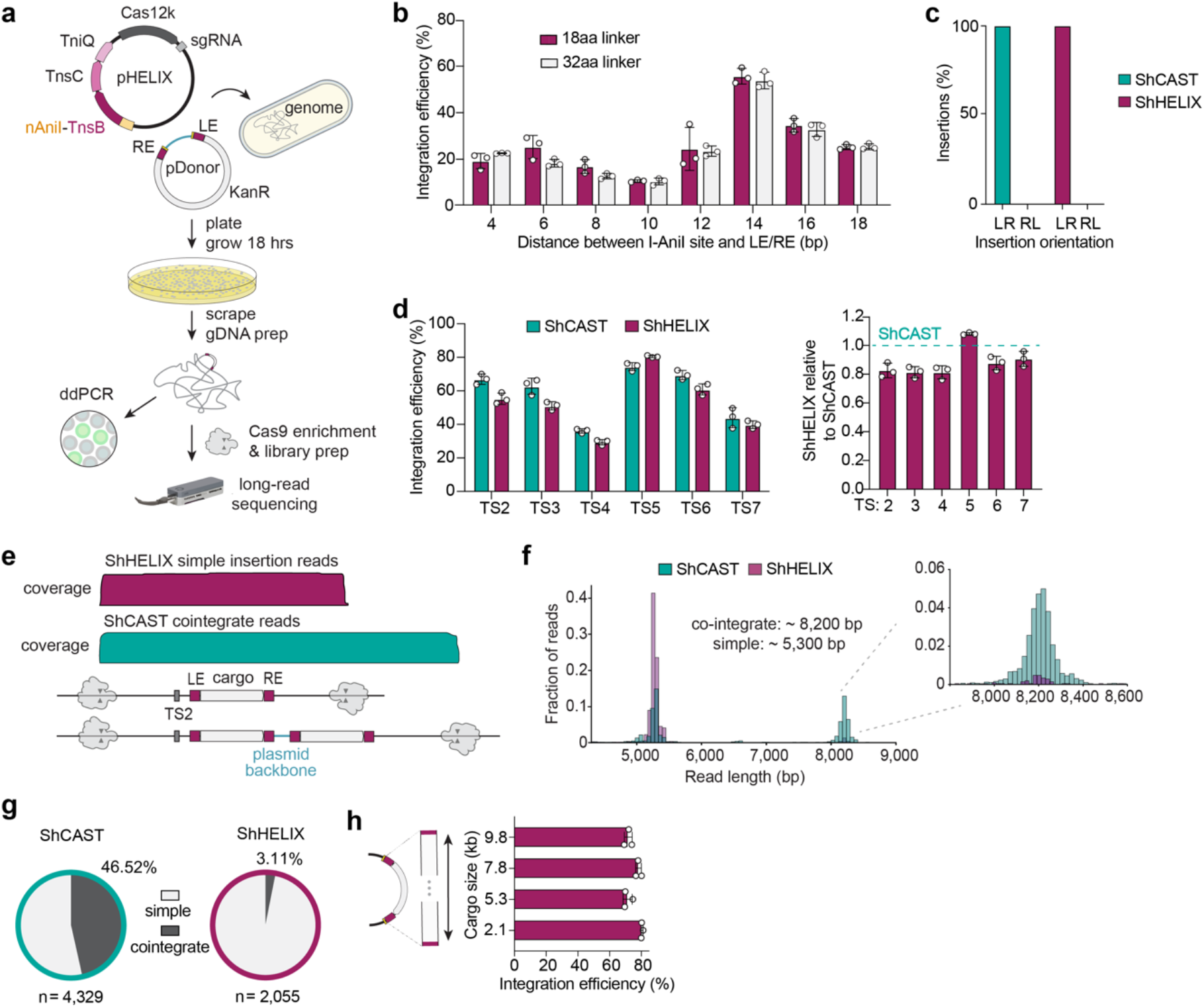
Characterization of DNA insertions on genomic targets using HELIX. **a**, Workflow for transposition experiments targeting the genome. **b**, Integration efficiencies when using two different amino acid linkers between nAniI and TnsB, an sgRNA against genomic target site 2 (TS2), and a set of eight donor plasmids with varying distances between the I-AniI sites and the LE/RE, as determined via ddPCR. **c**, Insertion orientation percentages when using ShCAST or ShHELIX targeting TS2 and using a pDonor with 14 bp spacing between the I-AniI site and the LE/RE **d**, Integration efficiencies across six genomic target sites for ShCAST and ShHELIX (left panel) and relative integration with ShHELIX normalized to ShCAST (right panel), assessed via ddPCR. **e**, Coverage of expected insertion products into the genome (TS2) from long-read sequencing using a subset of exemplary simple insertion reads for ShHELIX and co-integrate reads for ShCAST (coverage from ShHELIX co-integrate reads and ShCAST simple insertion reads omitted for simplicity). Transposed products were enriched prior to sequencing via Cas9 targeted enrichment. **f**, Read-length distribution of transposition products when using ShCAST and ShHELIX on genomic target site 2 (TS2) from long-read sequencing data. The top right panel is a zoomed in representation of the ∼8,200 bp read-length peak. **g**, Comparison of simple insertion and co-integrate product proportions at TS2 for ShCAST and ShHELIX, assessed via long-read sequencing. **h**, Integration efficiencies with ShHELIX and the sgRNA targeted to TS5, when using pDonors encoding cargoes of various sizes. Integration assessed via ddPCR. For **panels b, d**, and **h**, mean, SD, and individual data points shown for n = 3. LE and RE, left and right transposon ends, respectively; sgRNA, single guide RNA; ddPCR, droplet digital PCR.

Having identified an optimal I-AniI site to LE/RE spacing on pDonor for genome targeting, we then compared the integration efficiencies and product purities of ShCAST and ShHELIX across a range of genomic sites. ShHELIX retained robust RNA-guided targeted integration across six genomic target sites at levels comparable to ShCAST (**Fig. 2d**). To analyze the integration product purity of HELIX when targeting the genome at TS2, we performed long-read sequencing using a Cas9 targeted enrichment strategy^32^. Analysis of target-enriched reads when using ShCAST and ShHELIX that contained or lacked the cargo insertion showed that integration efficiencies calculated from our long-read sequencing data were similar to our ddPCR results at TS2 (**Sup. Fig. 5**). With ShCAST, we observed that 46.52% of insertion reads were co-integrates (**Figs. 2e-g**), which is generally lower than previously observed, albeit against a different target site and via alternate long-read sequencing methods^17^. With ShHELIX, we observed only 3.11% co-integrates, a 17-fold decrease compared to ShCAST (**Figs. 2e-g**).

Next, we assessed the ability of ShHELIX to integrate DNA cargos of various sizes. We performed transformations using donor plasmids harboring cargos of either a 5.2, 7.8, or 9.8 kb sequence (compared to pDonor with a 2.1 kb cargo used in previous experiments). For each transposition reaction using the larger cargos, ShHELIX showed comparably high efficiency of targeted DNA integration irrespective of cargo size (**Fig. 2h**). Together, our results demonstrate that ShHELIX is capable of highly active, unidirectional, cut-and-paste DNA insertions and is insensitive to cargo sizes up to at least 10 kb.

### Minimization of ShCAST and HELIX architectures

In contrast to the multi-component complexity of type I CASTs that require seven or eight protein subunits at varying stoichiometries, we wondered whether we could further simplify the typically 4 subunit type V-K CAST architecture. To do so, we designed 3-component systems by exploring fusions of various ShCAST subunits. We examined the fusion of one or two TniQ monomers to the N- or C-terminus of Cas12k, or the fusion of a single TnsC monomer on either Cas12k termini (**Fig. 3a**). Fusion of either TniQ or TnsC to the N-terminus of Cas12k substantially reduced integration efficiency (**Fig. 3b**). C-terminal fusions of TniQ to Cas12k retained up to 62.3% and 69.6% integration efficiency relative to that of unfused ShCAST, when one or two TniQ monomers were fused, respectively (**Fig. 3b** and **Sup. Fig. 6a**). We also observed that fusion of a TnsC monomer to the C-terminus of Cas12k retained relatively high integration efficiency (**Fig. 3b**). Given the proposed critical role of TnsC filament formation for transposition^30,33-35^, we speculate that filamentation may still occur, despite Cas12k fusion, or that only a single monomer is sufficient to enable transposition through a non-canonical mechanism (**Sup. Note 3**).

**Figure 3.**
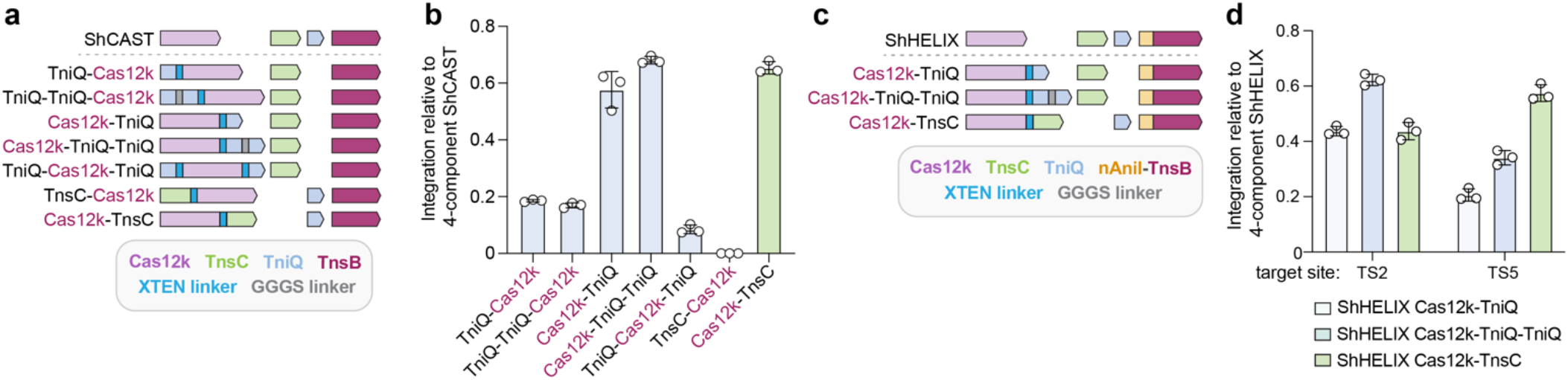
Integration efficiencies of 3-component CAST and HELIX systems. **a**, Schematic of fusion architectures to create 3-component ShCASTs. **b**, Integration efficiencies normalized to canonical, 4- component ShCAST for each 3-component ShCAST system, as determined by ddPCR (see **Sup. Fig. 6a** for absolute levels of integration). **c**, Schematic of 3-component ShHELIX systems. **d**, Integration efficiencies normalized to 4-component ShHELIX for each 3-component ShHELIX system (see **Sup. Fig. 6b** for absolute levels of integration). For **panels b** and **d**, mean, SD, and individual data points shown for n = 3.

Next, we determined whether analogous Cas12k fusions were similarly functional in the context of 3-component HELIX systems (**Fig. 3c**). We performed transformations using Cas12k-TniQ, Cas12k-TniQ- TniQ, or Cas12k-TnsC fusions along with the remaining HELIX modules and assessed integration efficiency. We observed RNA-guided integration between 22-36% at TS2 and 15-49% at TS5 depending on which fusion architecture was employed (**Fig. 3d** and **Sup. Fig. 6b**). Next-generation amplicon sequencing revealed that 4- and 3-component ShCAST and ShHELIX systems all exhibited the same PAM-to-LE insertion distance profile (**Sup. Fig. 7**), revealing that nAniI-TnsB fusions and TniQ or TnsC fusions to Cas12k, either separately or when combined, do not alter this property. These results demonstrate the feasibility of generating HELIX systems with fewer subunits through component fusions, and highlights protein engineering avenues to construct 3-component versions that exhibit comparable integration efficiencies.

### Extensibility of HELIX to type V-K CAST orthologs

All discovered type V-K CASTs lack TnsA^36^. This observation supports the evolutionary hypothesis that a Tn5053-like transposon, containing TnsB, TnsC, and TniQ, but not TnsA, co-opted and repurposed this CRISPR system^19^. Therefore, all type V-K CASTs would be expected to act through replicative transposition, leading to a substantial fraction of undesired co-integrate products. Thus, we explored HELIX as a generalizable approach to enable cut-and-paste DNA insertion with other diverse type V-K CASTs (**Fig. 4a**).

**Figure 4.**
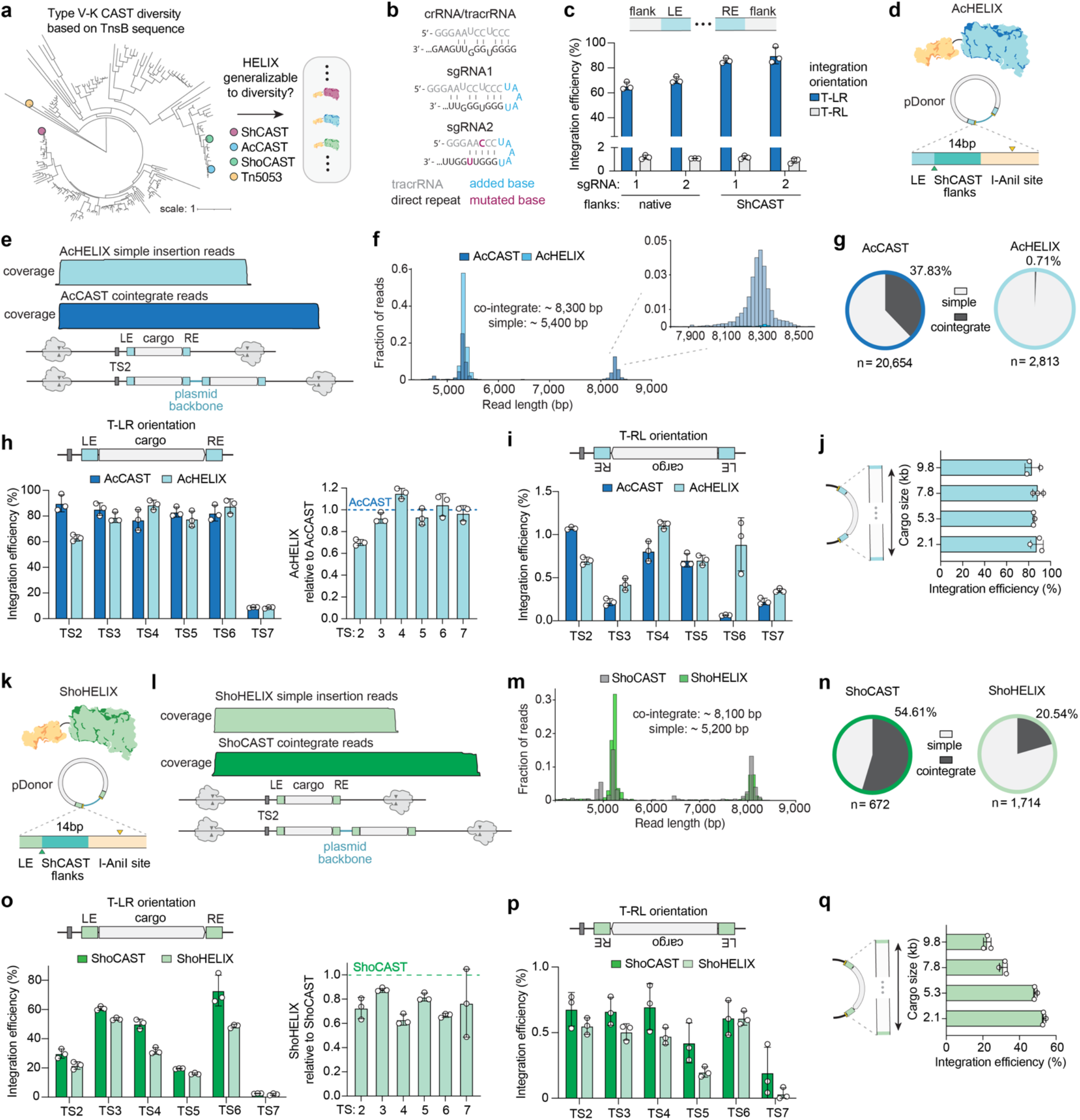
Extension of HELIX to type V-K CAST orthologs. **a**, Phylogenetic tree illustrating diversity of TnsB sequences from recently identified Type V-K CASTs^36^ CASTs used in the present study, as well as Tn5053, are noted. **b**, sgRNA designs for AcCAST. **c**, Integration efficiencies with AcCAST using two sgRNA designs (from **panel b**) and a donor plasmid with either native flanking sequence (as previously reported^14^) or ShCAST flanking sequence, assessed via ddPCR. **d**, Schematic of AcHELIX with 14 bp ShCAST flank sequence on pDonor. **e**, Coverage of insertion products into the genome (TS2) from long- read sequencing, displaying a selection of exemplary simple insertion reads for AcHELIX and co- integrate reads for AcCAST (coverage from AcHELIX co-integrate reads and AcCAST simple insertion reads omitted for simplicity). Transposed products were enriched prior to sequencing via Cas9 targeted enrichment. **f**, Read-length distribution of transposition products when using AcCAST and AcHELIX on TS2 from long-read sequencing data. The top right panel is a zoomed in representation of the ∼8.3 kb read-length peak. **g**, Comparison of simple insertion and co-integrate product proportions for AcCAST and AcHELIX, assessed via long-read sequencing. **h**,**i**, Integration efficiencies in the T-LR and T-RL orientations (**panels h** and **i**, respectively) across six genomic target sites for AcCAST and AcHELIX, assessed via ddPCR. In **panel h**, AcHELIX T-LR integration efficiency relative to AcCAST is shown in the right panel. All transformations contain the pDonor variant with ShCAST flanks and 14 bp spacing between the nAniI sites and LE/RE. **j**, Integration efficiencies when using AcHELIX using the sgRNA targeted to TS6 and pDonors encoding cargoes of various sizes, assessed via ddPCR. **k**, Schematic of ShoHELIX with 14 bp ShCAST flank sequence on pDonor. **l**, Coverage of expected insertion products into the genome (TS2) from long-read sequencing, displaying a selection of exemplary simple insertion reads for ShoHELIX and co-integrate reads for ShoCAST (coverage from ShoHELIX co-integrate reads and ShoCAST simple insertion reads omitted for simplicity). Transposed products were enriched prior to sequencing via Cas9 target enrichment. **m**, Read-length distribution when using ShoCAST and ShoHELIX on a genomic target (TS2) from long-read sequencing data. **n**, Comparison of simple insertion and co-integrate product proportions for ShoCAST and ShoHELIX, assessed via long-read sequencing. **o**,**p**, Integration efficiencies in the T-LR and T-RL orientations (**panels o** and **p**, respectively) across six genomic target sites for ShoCAST and ShoHELIX, assessed via ddPCR. **q**, Integration efficiencies when using ShoHELIX with a TS3-targeted sgRNA and pDonors encoding cargoes of various sizes, assessed via ddPCR. All ShoCAST and ShoHELIX transformations contain a pDonor variant with ShCAST flanks. For **panels c, h**-**j**, and **o**-**q**, mean, SD, and individual data points shown for n = 3. LE and RE, left and right transposon ends, respectively; sgRNA, single guide RNA.

To investigate the applicability of HELIX to other CAST orthologs, we characterized and optimized two previously reported type V-K CASTs from either *Anabaena cylindrica* (AcCAST) or a different strain of *Scytonema hofmannii* (ShoCAST). First, for the canonical AcCAST system, we designed two sgRNA scaffolds (**Fig. 4b**) and two pDonor architectures, the latter of which varied by containing different 25 bp sequences flanking the LE and RE (either as previously reported for AcCAST^14^ or using the ShCAST flanking sequences). With the two sgRNA designs that differed based on their crRNA-tracrRNA fusion points, we observed only a modest difference in integration efficiency (**Figs. 4b** and **4c**). However, the pDonor containing ShCAST flanking sequences resulted in increased absolute integration efficiencies of 19.6% or 20.4% for sgRNA-1 and sgRNA-2, respectively (1.28- and 1.31-fold increases over pDonor with the native AcCAST flanks; **Fig. 4c**). As we previously observed for ShCAST (**Sup. Fig. 1b**), these results suggest that the sequences directly adjacent to the LE and RE on pDonor are an important determinant of type V-K CAST-mediated integration efficiency. Considering the flank sequences, as well as other potential determinants of integration efficiency, such as pDonor copy number (**Sup. Fig. 8**), are opportunities to optimize diverse CASTs. Additionally, AcCAST showed a minimal, though still detectable, number of T-RL oriented insertions, making it a near-complete unidirectional inserter (**Fig. 4b**).

We constructed AcHELIX comprising a nAniI-TnsB fusion along with the sgRNA-2 design and a pDonor harboring I-AniI sites 14 bp from the LE/RE separated by ShCAST flanking sequence (**Fig. 4d**). To determine the integration product purity with AcHELIX compared to AcCAST when targeting the genome, we performed long-read sequencing following Cas9 target enrichment (**Fig. 4e**). While with AcCAST we observed 37.83% co-integrate products, for AcHELIX we found only 0.71%, representing a 53-fold improvement in product purity with AcHELIX (**Figs. 4f** and **4g**). Across six genomic targets, AcHELIX retained comparable RNA-guided DNA integration and insertion directionality to AcCAST (**Figs. 4h** and **4i**). Additionally, similar to ShHELIX, AcHELIX demonstrated no decrement in efficiency when integrating cargo sequences of various sizes up to 9.8 kb, maintaining over 83% integration efficiency for all four cargo sizes at TS6 (**Fig. 4j**). Thus, similar to ShHELIX, AcHELIX is an efficacious engineered CAST with near-perfect simple insertion product purity for DNA insertions of various sizes.

Next, we characterized ShoCAST and ShoHELIX utilizing a pDonor with a 14 bp spacing separating the I-AniI site and LE/RE with ShCAST flanking sequence (**Fig. 4k**). We performed genome-targeting experiments with ShoCAST and ShoHELIX using a previously reported sgRNA^16^ against TS2. Characterization of the insertion products via long-read sequencing revealed 54.61% co-integrates for ShoCAST and 20.54% for ShoHELIX, demonstrating a 2.6-fold reduction in co-integrates when using ShoHELIX (**Figs. 4l-4m**). Across genomic targets TS2-TS7, we observed a range of integration efficiencies, with ShoHELIX exhibiting comparable integration to ShoCAST (**Fig. 4o**). Similar to AcCAST and AcHELIX, the directionality of ShoCAST and ShoHELIX insertions were predominantly in the T-LR orientation, albeit with minimal but detectable T-RL insertions (**Fig. 4o** and **4p**). Additionally, in contrast to ShHELIX and AcHELIX, ShoHELIX showed a decrease in integration efficiency with increasing cargo size on pDonor at TS3 (**Fig. 4q**).

## Discussion

CASTs are an emergent class of genome editing technologies that enable programmable DNA insertions without reliance on recombination, sequence-specific recombinases, or DSBs. However, the currently discovered and characterized type I and V-K systems have limitations that restrict their ease of use, including size (**Sup. Fig. 9**), stoichiometric and component complexity, and/or insertion product purity. To overcome these constraints, we developed HELIX, which harnesses the technological advantages of type V-K CASTs, and employs a nHE fusion and a modified donor plasmid to achieve programmable and efficient cut-and-paste DNA insertion similar to type I CASTs. We show that HELIX dramatically increases simple insertion product purity on plasmid and genomic targets in *E. coli* and retains robust RNA-guided transposition at or near wild-type levels. Additionally, we show the feasibility of simplifying CAST and HELIX systems to 3-component systems via subunit fusions to Cas12k, the utility of which will be improved by future engineering efforts to increase their integration efficiencies. Together, our approaches are the first descriptions of CAST engineering and highlight how other naturally occurring enzymes can be leveraged to augment CAST properties.

We demonstrate that HELIX is efficacious across three type V-K CAST orthologs, establishing the universality of this approach. The continued discovery of CASTs will not only identify new systems with useful characteristics but will also create a platform to develop next-generation HELIX technologies with diverse and useful properties. The ongoing metagenomic discovery of CASTs has identified a variety of transposon architectures with a range of properties, including compact Type V-K CASTs that may prove useful for genome editing applications^36^. Optimization of HELIX properties for each CAST ortholog (e.g. amino acid linkers, spacings between nHE sites and LE/RE, nHE selection, donor flanking sequences, etc.) will lead to higher transposition efficiencies while leveraging the unique properties of each CAST.

Amidst our CAST characterization, we also discovered that the flanking sequencing directly adjacent to the LE/RE on pDonor can influence integration efficiency. This finding suggests that future efforts to better understand and augment the flanking sequences, and potentially other parameters of CASTs, holds promise to further enhance transposition efficiencies. A more complete understanding of the determinants of CAST and HELIX integration efficiencies and product purities (e.g. donor flanking sequences, target site selection, cargo size, more comprehensive definition of PAM, etc.), as well as specificity, will be crucial in developing and tuning these systems.

In summary, HELIX is a generalizable approach that enables programmable, cut-and-paste, unidirectional, minimal component, and recombination-independent DNA insertions without DSBs. Efforts to create high specificity and hyperactive variants of type V-K CASTs, potentially through TnsC and Cas12k or TnsB and Cas12k engineering, respectively, could benefit next-generation HELIX systems. In parallel, HELIX holds potential to extend the use of CASTs to various organisms, given its improved subunit, stoichiometric, and insertion mechanism simplicity. Overall, HELIX demonstrates the prospect of applying engineering approaches to optimize and overcome the limitations of CASTs, with implications for the development of optimal DNA insertion technologies.

### Contributions

C.J.T conceived of the idea, designed and performed experiments, and wrote the manuscript draft. B.O. performed experiments for ShoHELIX and cargo size comparisons. B.P.K contributed to experimental design and supervised the study. C.J.T and B.P.K revised the manuscript; all authors approved the manuscript.

## Acknowledgments

We thank L.T. Hille, C.R.R. Alves, and R.A. Silverstein for suggestions about the manuscript, P.M. Boone for assistance with ddPCR, and B.L. Stoddard for advice. C.J.T. was supported by a National Science Foundation Graduate Research Fellowship Grant No. 2020295403. B.P.K was supported by a Mass General Hospital Howard M. Goodman Fellowship.

## Data Availability

All sequencing data will be deposited with the NCBI Sequence Read Archive (SRA).

## Methods

### Plasmids and oligonucleotides

All plasmids used in this study and selected sequences are listed in **Supplementary Table 1**. New plasmids were generated via isothermal assembly or Golden Gate assembly, some of which have been deposited with Addgene (**Supplementary Table 1**). pHelper and pDonor plasmids for ShCAST and AcCAST, as well as pTarget, were gifts from Feng Zhang (Addgene plasmid numbers 127921, 127924, 127923, 127925, 127926). For gRNA-encoding plasmids, spacer sequences were cloned into pCAST and pHELIX plasmids via Golden Gate assembly with SapI (New England Biolabs, NEB). Target site features for all gRNAs used in this study are found in **Supplementary Table 2**. Oligonucleotides and probes used in this study were purchased from Integrated DNA Technologies (IDT) and are listed in **Supplementary Table 3**. Gene fragments for construct cloning were ordered from Twist Biosciences; synthetic SpCas9 sgRNAs were ordered from Synthego (**Supplementary Table 2**).

### Transposition reactions targeting plasmids and genomic sites

Transformations for plasmid targeting experiments were performed in chemically competent PIR1 cells containing pTarget (original PIR1 strain obtained from Invitrogen), using 25 ng of pCAST or pHELIX and 25 ng of pDonor. Transformed cells were recovered for 1 hr at 37 °C in S.O.C. and then plated on LB agar plates containing 50 μg/mL kanamycin, 25 μg/mL chloramphenicol, and 100 μg/mL carbenicillin. Plates were incubated at 37 °C for 18 hrs. Colonies were counted, scraped, and plasmid DNA extracted via miniprep (Qiagen). The resulting plasmid pool was used for downstream analysis via junction PCR and long-read sequencing. Junction PCRs were analyzed via QIAxcel Capillary Electrophoresis (Qiagen) and visualized with QIAxcel ScreenGel Software (v1.5.0.16; Qiagen).

Transformations for genome targeting experiments were performed using PIR1 cells (or PIR2 cells (Invitrogen) for **Sup. Fig. 7**) and 25 ng of pCAST or pHELIX and 25 ng of pDonor. Transformed cells were recovered for 1 hr at 37 °C in S.O.C. and then plated on LB agar plates containing 50 μg/mL kanamycin and 100 μg/mL carbenicillin. For transformations including ShCAST, ShHELIX, ShoCAST, or ShoHELIX plasmids, plates were incubated at 37 °C for 18 hours; for AcCAST and AcHELIX transformations, plates were incubated at 37 °C for 24 hrs due to comparatively smaller colonies (though approximately the same in number). Colonies were scraped and gDNA was harvested using Wizard Genomic DNA Purification Kit (Promega) for downstream analysis via ddPCR and long-read sequencing.

### Assessment of integration efficiency via ddPCR

Plasmid or genomic DNA was first normalized to 10 ng/μL or 100 ng/μL, respectively, and then further diluted to 0.2 ng/μL or 2 ng/μL working stocks, respectively, for ddPCR assays. Insertion events were measured using target-specific primers and a donor-specific probe (**Supplementary Table 3**). ddPCR reactions contained 20 pg of plasmid DNA or 2 ng gDNA, 250 nM each primer, 900 nM probe, and ddPCR supermix for probes (no dUTP) (BioRad) in 20 μL reactions, and droplets were generated using a QX200 Automated Droplet Generator (BioRad). Thermal cycling conditions were: 1 cycle of (95 °C for 10 min), 40 cycles of (94 °C for 30 sec, 58 °C for 1 min), 1 cycle of (98 °C for 10 min), hold at 4 °C. PCR products were analyzed using a QX200 Droplet Reader (BioRad) and absolute quantification of inserts was determined using QuantaSoft (v1.7.4). Total template DNA was also analyzed, and integration efficiencies were calculated by inserts/template*100.

### Long-read sequencing of plasmid and genomic integrations

Integration product purity was analyzed via long-read sequencing using the plasmids resulting from plasmid targeting transposition reactions. Transposed products were enriched by electroporating approximately 100 ng of plasmid pool into Endura Electrocompetent Cells (Lucigen), which are a non-PIR strain that limits recombination. Cells were recovered for 1 hr at 37 °C in S.O.C. and spread on LB agar plates containing 50 μg/mL kanamycin and 25 μg/mL chloramphenicol. Plates were incubated at 30 °C (to limit recombination) for 24 hrs, scraped, and plasmid DNA extracted via miniprep. Enriched plasmids were digested with EcoRV (NEB) for 8 hrs at 37 °C. Amplification-free long-read sequencing library preparation (Oxford Nanopore Technologies, SQK-LSK109) was performed using a barcode expansion kit (Oxford Nanopore Technologies, NBD-104). The final pooled library was loaded onto an R9.4.1 flow cell and sequenced for 24 hrs.

To conduct long-read sequencing of genome-targeted insertions, we performed an amplification-free Cas9 targeted enrichment protocol to improve sequencing selectively of the intended on-target sites (Oxford Nanopore Technologies, SQK-CS9109; sgRNAs listed in **Supplementary Table 2**). The resulting library was loaded onto an R9.4.1 flow cell and sequenced for 30 hrs.

### Data processing of long-read sequencing results

Fast5 files were base called in real time using Miknow (v21.06.9) with the fast base calling model, and the resulting FastQ files were filtered for Q score > 8. BBDuk from the BBTools suite^37^ was used to filter for reads containing 20bp of LE, RE, and the target site spacer sequence with a maximum hamming distance of 2. Of these reads, those containing a 20 bp sequence (with a maximum hamming distance of 2) found in the plasmid backbone (not expected to occur in simple insertion products) were categorized as potential co-integrates and those not containing this sequence were categorized as potential simple insertions. Reads for plasmid-targeting experiments were additionally filtered for appropriate read length. Reads containing products assigned as simple insertions or co-integrates were merged into a single FastQ file and aligned to either a synthetic simple insertion or co-integrate product with Minimap2^38^ specified with the *map-ont* parameter. Coverage plots were generated from an exemplary set of 100 reads using Geneious (v2021.2.2) and its inbuilt aligner (medium sensitivity and an iteration of up to 5 times). Sam files containing aligned reads were also produced and used to generate length histograms.

### Analysis of insertion distance using targeted sequencing

PAM-to-LE insertion distances were assessed by next-generation sequencing using a 2-step PCR-based library construction method. 50 ng of genomic DNA from genome-targeting experiments were PCR amplified with 30-cycles using Q5 High-fidelity DNA Polymerase (NEB) and primers which bind just outside of TS2 or just inside of LE. PCR products were purified using paramagnetic beads prepared as previously described^39,40^. 20 ng of purified PCR product was used as template for a second, 10-cycle PCR to add Illumina barcodes and adapter sequences. PCR products were purified prior to quantification via QuantiFluor (Promega) and combined into an equimolar pool. Final libraries were quantified by qPCR (KAPA Library Quantification Kit; Roche 7960140001) and sequenced on a MiSeq using a 300- cycle v2 kit (Illumina).

### Data processing of targeted sequencing results

Paired FastQ reads were first filtered for Q>30 using BBDuk from the BBTools suite and merged via BBMerge. Reads containing 20bp of TS2 and 20bp of the terminal LE, each with a maximum hamming distance of 1, were then extracted. Each read was then trimmed of the sequence upstream of and including the PAM and downstream of and including the LE, resulting in only the sequence between the PAM and LE (i.e. site of insertion). Lengths of the resulting reads were calculated and used to plot PAM-to-LE insertion distance profiles.

