## Supplementary Materials for "Cut-and-Paste DNA Insertion with Engineered Type V-K CRISPR-associated Transposases"

Supplementary Tables

**Supplementary Table 1**: Plasmids and sequences in this study

**Supplementary Table 2**: gRNA spacer sequences used in this study

**Supplementary Table 3**: Oligonucleotides and probes used in this study

***Note:*** *All Supplementary Tables are attached together as a single .xlsx file, separated by tabs*.

Supplementary Notes

**Supplementary Note 1**: Expanded discussion of Y2 ShHELIX results p.2

**Supplementary Note 2**: ShHELIX control experiments p.2

**Supplementary Note 3**: Mechanistic implications of Cas12k-TnsC fusions p.3

Supplementary Figures and Legends

**Supplementary Figures 1-8** p.5-13

Supplementary References

**Supplementary References** p.14

### Supplementary Notes:

**Supplementary Note 1: Expanded discussion of Y2 ShHELIX results**

While developing and characterizing ShHELIX, we also assessed whether the Y2 nAniI variant, shown to have a 9-fold higher affinity for its cognate target site[1], would enable a further increase in simple insertion product purity. With the Y2 ShHELIX construct, we observed a stark decrease in transformant colonies (**Sup. Fig. 3a**), as compared to ShCAST or non-Y2 ShHELIX (**Sup. Fig. 1a**). Moreover, this decrease varied with the spacing between the I-AniI site and LE/RE on pDonor, where a 14 bp spacing showed the highest number of colony-forming units (CFUs) (also aligning with the spacing giving the highest integration efficiency via ddPCR on plasmid and genomic targets). In combination with a similar observation when using a Lib4 I-AniI site (as shown in **Fig. 1k**), where the Lib4 I-AniI site was previously shown to increase wild type I-AniI affinity site by 5-fold[2], we recognized a correlation between the affinity of I-AniI for its target sequence and the number of colonies present on plates selecting for pShHELIX or pShCAST, pDonor and/or transposed product, and pTarget.

While further studies into the mechanism of HELIX will fully elucidate the cause of this observation, we speculate that a combination of two phenomena may be occurring. First, the higher affinity of Y2 nAniI for its target, or when using nAniI with a Lib4 site, leads to an increased prevalence of DNA double-strand breaks (DSBs) on pDonor at early time points in the post-transformation recovery. In the absence of rapid and efficient cargo integration into pTarget, the AniI-caused DSBs result in a loss of Kanamycin resistance due to pDonor degradation prior to transposition. In this scenario, colony counts for different spacings on pDonor may correlate with higher or lower integration efficiencies. For example, for spacings where transposition is most efficient and rapid, the loss in CFUs is less striking because integration into pTarget occurs more rapidly than DSBs on pDonor. A second hypothesis is that the higher affinity of Y2 nAniI for its target, or when using nAniI with a Lib4 site, leads to an increased occurrence of DSBs on pDonor. Given the high copy number of pDonor in PIR1 cells, this could result in SOS response induction and cell death.

**Supplementary Note 2: ShHELIX control experiments**

While performing long-read sequencing of transposition products resulting from plasmid-targeting experiments, we included several control conditions. Firstly, we performed experiments using a catalytically attenuated I-AniI variant (harboring K227M and Q171K mutations[3]) to create a 'dead' ShHELIX (dShHELIX). With dShHELIX, we observed a 1.8-fold decrease in co-integrate products compared to wild-type ShCAST (**Sup. Fig. 4a** and **Fig. 1i**, respectively). We hypothesize that this somewhat unexpected decrease in co-integrate products is the result of incomplete inactivation of I-AniI catalysis, which might lead to low-level 5' pDonor nicking (at a rate slower than nAniI-based ShHELIX). Indeed, the I-AniI Q171K variant has previously been shown to exhibit residual nicking activity on both DNA strands *in vitro*[3].

Secondly, we performed experiments using a pDonor variant that does not harbor I-AniI sites. In transformations with ShHELIX and this modified pDonor lacking I-AniI sites, we observed a 1.7-fold decrease in co-integrates relative to ShCAST (**Sup. Fig. 4a** and **Fig. 1i**, respectively). We hypothesize that this could be due to low-level I-AniI activity on sequences flanking the LE and RE. A previous study which mutated each base in the I-AniI recognition sequence to all other bases revealed that specificity of nAniI is greatest across base pair positions ±3, 4, 5, and 6 in each half-site and least specific across bases -2 to +1 and bases at the outer edges of the recognition sequence[3]. From this data, a minimal approximate core sequence of 5'-GAGGNNNCTCTG-3' is necessary for I-AniI recognition, with decreased activity depending on the base substituted. While we could not identify an exact sequence match, we note that sequences similar to these core motifs occur on pDonor at 5'-GTGGNNNNGTCTA-3' (11 bp from the LE) and 5'-GAGGNNNCATTG-3' (13 bp from the RE), the latter being in an orientation that would give a nick on the same strand as TnsB (see next point). Low-level nicking on these flanking sequences at these degenerate I-AniI core sequences might lead to a slight increase in simple insertion product purity (as observed).

Thirdly, we performed experiments using 'flipped' I-AniI sites on pDonor oriented to confer a nick on the same strand as TnsB. In experiments using a flipped I-AniI site pDonor, we observed a 10-fold decrease in co-integrates with ShHELIX relative to ShCAST (**Sup. Fig. 4b**). We hypothesize that this reduction in co-integrates might be the result of an alternative transposition mechanism involving hairpin intermediates and processing (MuA of the Mu transposon has been shown to successfully process hairpins prior to insertion[4]) or possibly via Shapiro intermediate cleavage (**Sup. Fig. 4c**).

### <u>Supplementary Note 3</u>: Mechanistic implications of Cas12k-TnsC fusions

Recent structural studies have provided insight into the mechanism of ShCAST-mediated DNA insertion[5,6]. These studies suggest that TnsB recruitment to TniQ-capped TnsC filaments simulates filament disassembly, exposing the target site and inducing insertion at a coordinated distance from the sgRNA-Cas12k-DNA complex. Our experiments with fusions of Cas12k to a TnsC monomer in the context of ShCAST or ShHELIX (**Fig. 3**) are interesting given these proposed mechanisms, particularly regarding the role of TnsC filamentation in recruiting downstream transposition machinery. Additionally, since the extent of TnsC filament disassembly (or the footprint of TniQ alone or bound to TnsC) may define the insertion distance from bound DNA-bound Cas12k for canonical 4-component ShCAST, it is interesting that our Cas12k-TnsC fusions (in the context of ShCAST and ShHELIX systems) enable targeted DNA insertion with the same insertion distance profiles as the canonical 4-component ShCAST and ShHELIX systems (**Sup Fig. 7**). We speculate that TnsC filamentation may still occur, despite Cas12k fusion, or that only a single TnsC subunit fused to Cas12k is sufficient to enable transposition. In the latter case, it is possible that TnsB-mediated depolymerization collapses TnsC filaments to a single monomer, which results in the fixed insertion distance profile observed for natural systems and would align with the identical profile observed for our monomer fusion. Alternatively, TnsC may not be involved in insertion distance determination, and a TniQ-defined insertion distance model may be more plausible. Our results with Cas12k-TnsC fusions provide unexpected insight into the role of TnsC in ShCAST-mediated transposition, motivating further studies to elucidate the transposition mechanism of both natural CAST and engineered HELIX 3- or 4-component systems.

### Supplementary Figures and Legends:

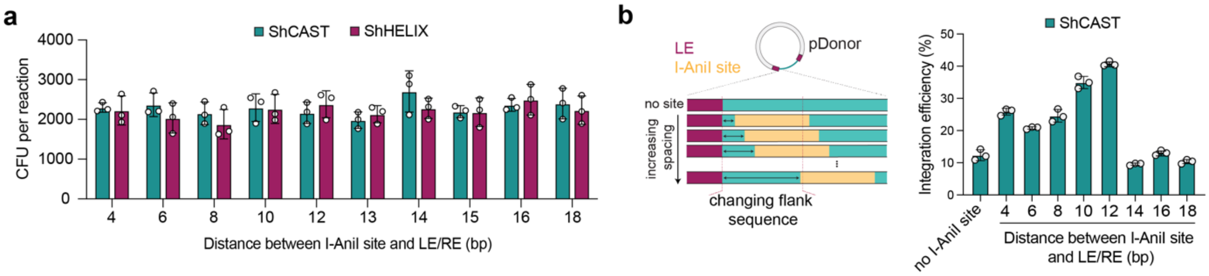


**Supplementary Figure 1. Characterization of plasmid-targeting experiments. a**, Colony-forming units (CFUs) from transformations with ShCAST and ShHELIX plasmids when using a series of pDonor plasmids bearing various spacings between the I-AniI sites and LE/RE. **b,** Schematic of donors bearing modified flank sequences (left panel), and integration efficiencies when using ShCAST and an array of pDonors with different LE/RE flank sequences (corresponding to the ShHELIX pDonors bearing different spacings between the I-AniI sites and the LE/RE), assessed via ddPCR (right panel). For **panels a** and **b**, mean, SD, and individual data points shown for *n* = 3. LE and RE, left and right transposon ends, respectively.

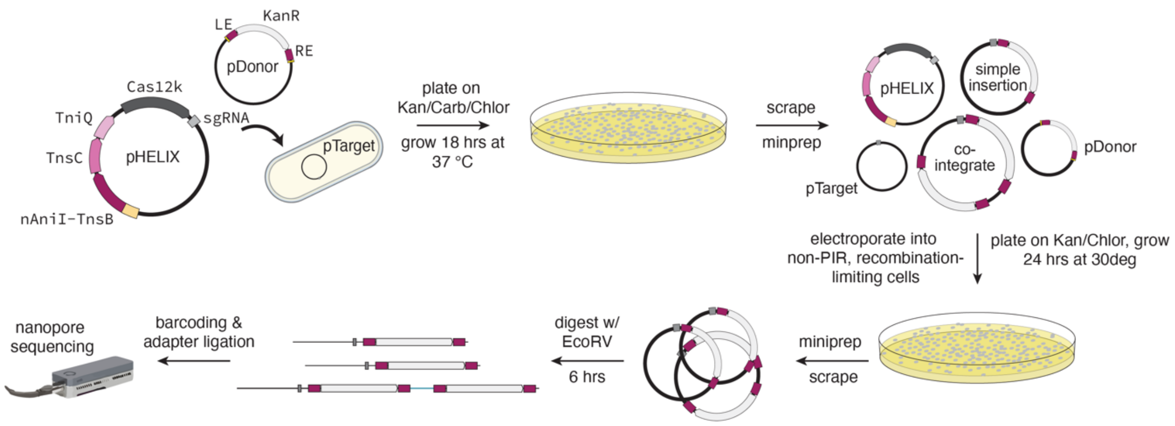


**Supplementary Figure 2. Workflow for plasmid enrichment prior to long-read sequencing.** Schematic of the protocol to enrich for transposed plasmid products to improve read-depth of intended products via long-read sequencing. sgRNA, single guide RNA; LE and RE, left and right transposon ends, respectively.

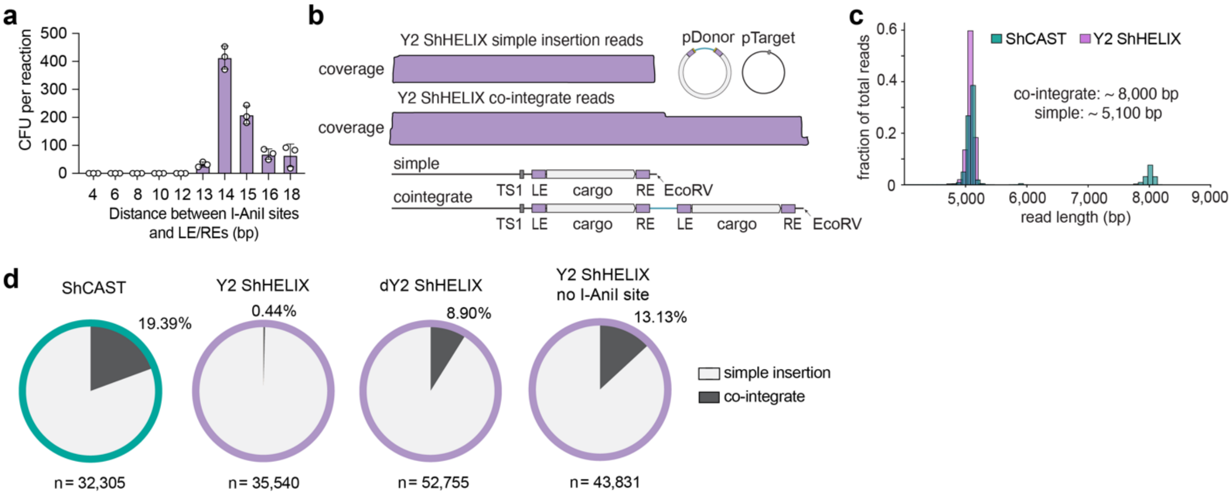


**Supplementary Figure 3. Characterization of Y2 ShHELIX. a,** Colony-forming units (CFUs) from transformations with Y2 ShHELIX plasmids when using a series of pDonor plasmids bearing various spacings between the I-AniI sites and LE/RE. **b,** Coverage of expected insertion products into pTarget from long-read sequencing, displaying an exemplary subset simple insertion or cointegrate reads for Y2 ShHELIX. **c**, Read length distribution when using ShCAST and Y2 ShHELIX with a sgRNA targeting TS1 on pTarget. **d,** Comparison of simple insertion and co-integrate product proportions via long-read sequencing for various conditions using Y2-ShHELIX. For **panel a**, mean, SD, and individual data points shown for *n* = 3. LE and RE, left and right transposon ends, respectively.

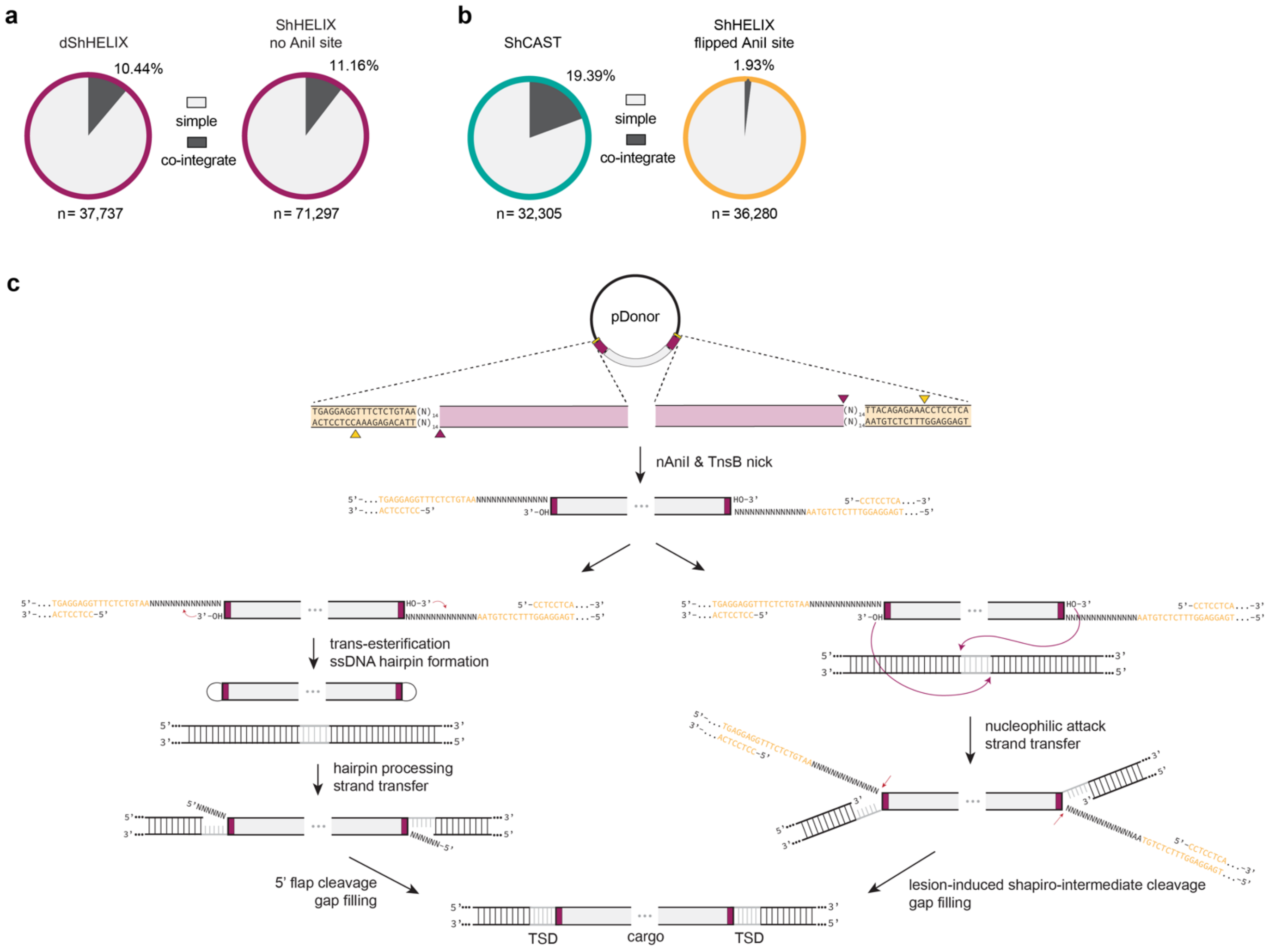


**Supplementary Figure 4. ShHELIX control experiments. a,** Comparison of simple insertion and co-integrate product proportions via long-read sequencing for a HELIX variant with a catalytically attenuated nAniI (dShHELIX) and when using HELIX with a pDonor without I-AniI sites. **b,** Comparison of simple insertion and co-integrate product proportions via long-read sequencing for ShCAST and ShHELIX when using a pDonor with flipped I-AniI sites that place the nAniI nicking sites on the same strand as the nick from TnsB. **c,** Potential alternative mechanisms enabling cut-and-paste DNA insertion with a pDonor containing a flipped I-AniI site. TSD, target site duplication.

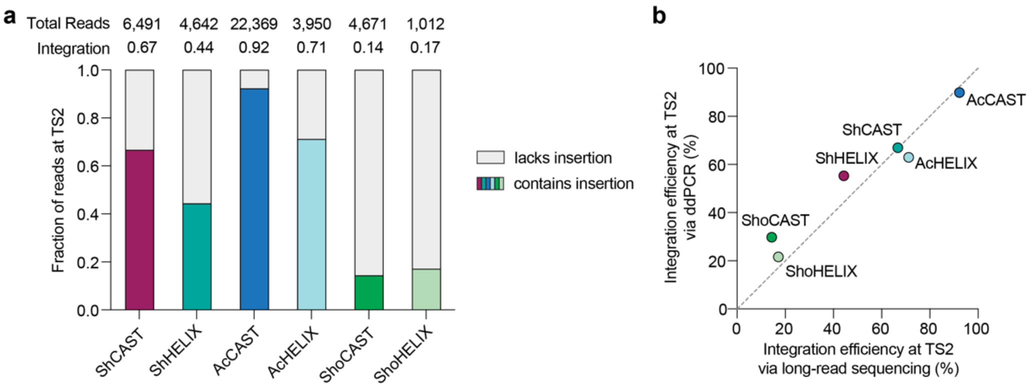


**Supplementary Figure 5. Integration efficiency based on long-read sequencing. a,** Integration efficiencies at TS2 when using CAST and HELIX systems, assessed via long-read sequencing. Stacked bars represent the fraction of Cas9-enriched target reads that lack or contain the cargo insertion. Integration (colored portion of each bar) represents the number of reads that contain the cargo insertion divided by the total number of targeted reads. **b,** Comparison of integration efficiencies for each system as measured by ddPCR or by Cas9-enriched long-read sequencing. The dashed grey line denotes the diagonal (agreement between the two types of measurements).

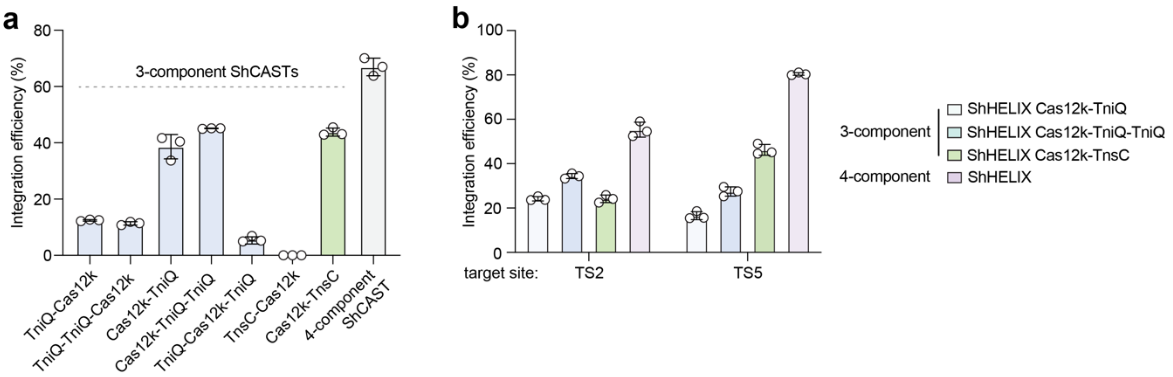


**Supplementary Figure 6. Absolute integration efficiencies for more minimal CAST and HELIX systems. a,b,** absolute integration efficiencies when targeting the genome at TS2 for 3- and 4-component ShCASTs (**panel a**), and when targeting TS2 or TS5 for 3- and 4-component ShHELIX systems (**panel b**). Efficiencies were assessed via ddPCR and used to calculate relative integration as shown in **Fig. 3**. For **both panels**, mean, SD, and individual data points shown for $n$ = 3.

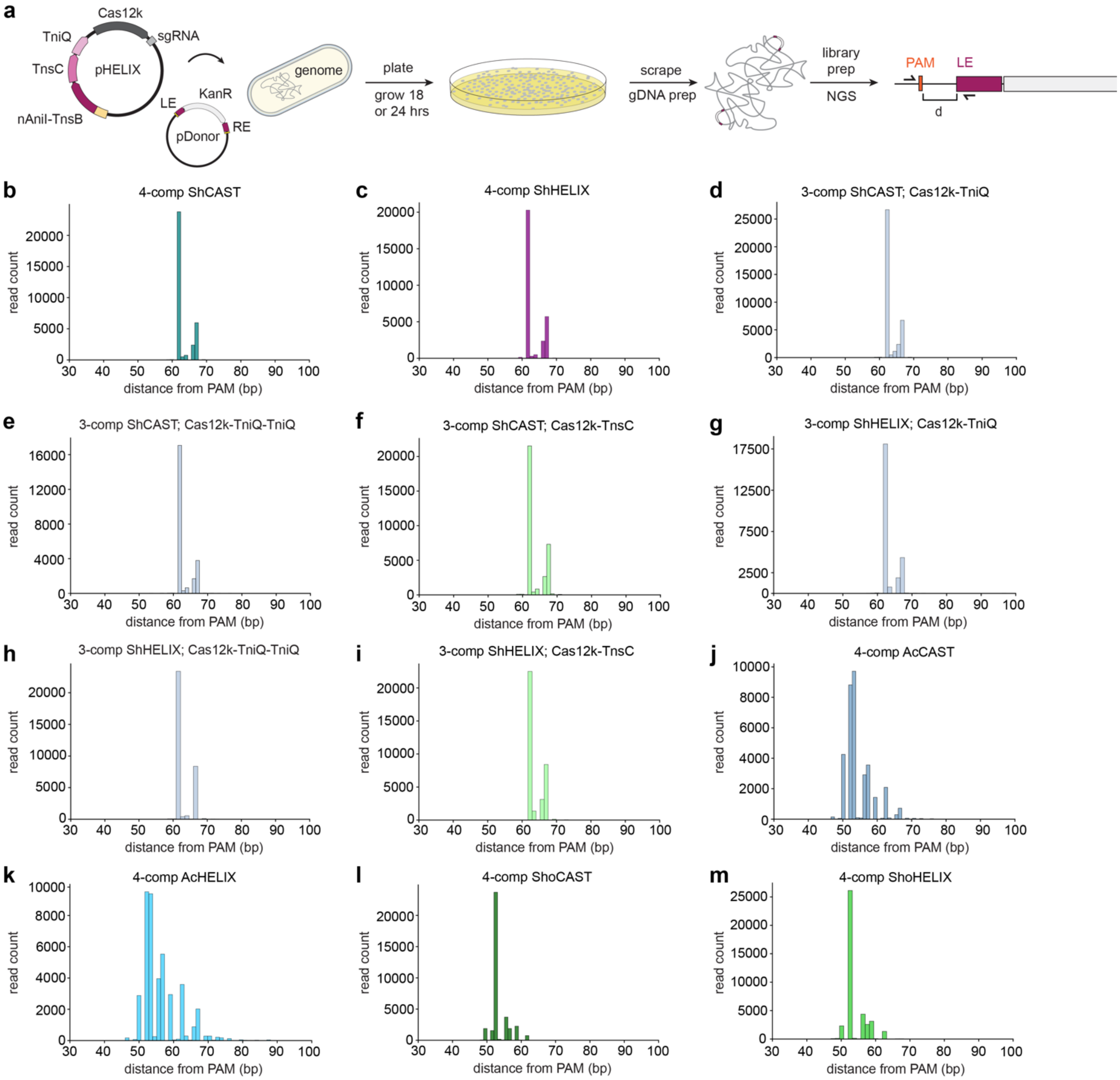


**Supplementary Figure 7. Cargo insertion distance from the PAM. a,** Schematic of the workflow to characterize PAM-to-LE insertion distances via next-generation targeted sequencing. PAM-to-LE insertion distance profiles for various CAST and HELIX constructs shown in **panels**: **b**, ShCAST (4-components); **c,** ShHELIX (4-components); **d,** ShCAST with Cas12k-TniQ (3-components); **e,** ShCAST with Cas12k-TniQ-TniQ (3-components); **f,** ShCAST with Cas12k-TnsC (3-components); **g**, ShHELIX with Cas12k-TniQ (3-components); **h,** ShHELIX with Cas12k-TniQ-TniQ (3-components); **i,** ShHELIX with Cas12k-TnsC (3-components); **j,** AcCAST (4-components); **k,** AcHELIX (4-components); **l,** ShoCAST (4-components); **m,** ShoHELIX (4-components). sgRNA, single guide RNA; PAM, protospacer adjacent motif; LE and RE, left and right transposon ends, respectively; NGS, next-generation sequencing.

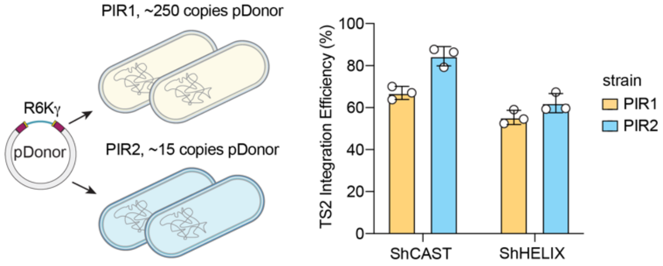


**Supplementary Figure 8. Influence of pDonor copy number on integration efficiency.** Integration efficiencies using ShCAST and ShHELIX and an sgRNA targeting genomic TS2 in two different bacterial strains that modulate pDonor copy number (where PIR1 and PIR2 cells maintain pDonor at approximately 250 and 15 copies, respectively). Integration efficiencies assessed via ddPCR; mean, SD, and individual data points shown for $n$ = 3. R6Kγ, origin of replication that requires the gene, *PIR,* to replicate.

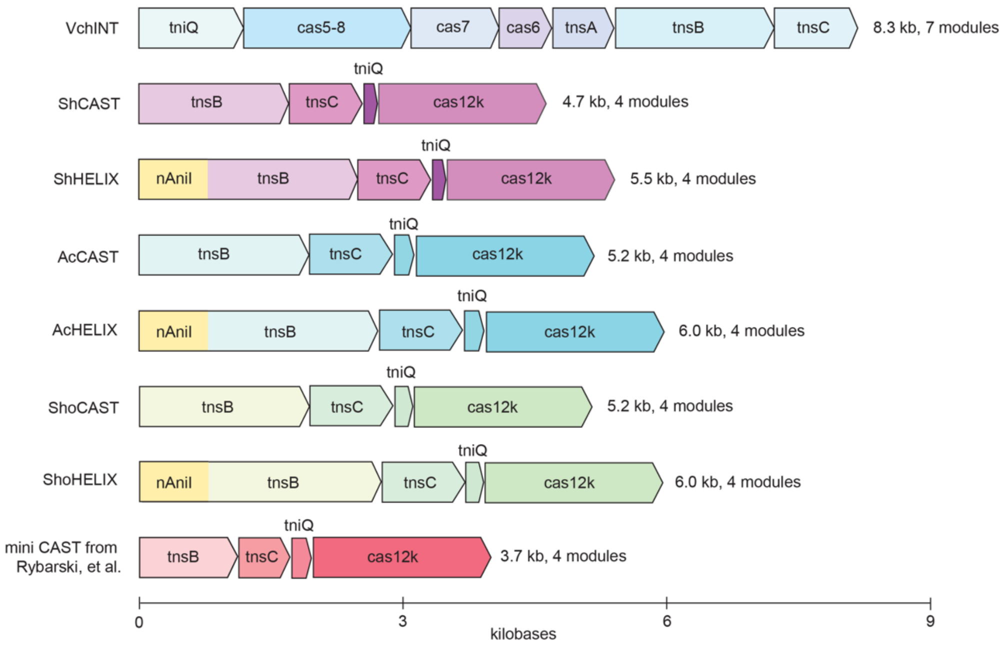


**Supplementary Figure 9. Coding sequence and component number comparison of CAST and HELIX systems**. Coding sizes and number of protein subunits for prototypical type I CASTs, type V-K CASTs, and HELIX systems for those in this study, as well as a recently described mini CAST from metagenomic mining[7]. nAniI, nicking I-AniI (K227M).
